# Brain tissue properties and morphometry assessment after chronic complete spinal cord injury

**DOI:** 10.1101/547620

**Authors:** Andreas Hug, Adriano Bernini, Haili Wang, Antoine Lutti, Johann M.E. Jende, Markus Böttinger, Marc-André Weber, Norbert Weidner, Simone Lang

## Abstract

There is much controversy about the potential impact of spinal cord injury (SCI) on brain’s anatomy and function, which is mirrored in the substantial divergence of findings between animal models and human imaging studies. Given recent advances in quantitative magnetic resonance imaging (MRI) we sought to tackle the unresolved question about the link between the presumed injury associated volume differences and underlying brain tissue property changes in a cohort of chronic complete SCI patients. Using the established computational anatomy methods of voxel-based morphometry (VBM) and voxel-based quantification (VBQ) we performed statistical analyses of grey matter volume and parameter maps indicative for brain’s myelin, iron and free tissue water content in complete SCI patients (n=14) and healthy individuals (n=14). Our whole-brain analysis showed significant white matter volume loss in the rostral and dorsal part of the spinal cord consistent with Wallerian degeneration of proprioceptive axons in the lemniscal tract in SCI subjects, which correlated with spinal cord atrophy assessed with quantification of the spinal cord cross-sectional area at cervical level. The latter finding suggests that Wallerian degeneration of the lemniscal tract represents a main contributor to the observed spinal cord atrophy, which is highly consistent with preclinical ultrastructural/histological evidence of remote changes in the central nervous system secondary to SCI. Structural changes in the brain representing remote changes in the course of chronic SCI could not be confirmed with conventional VBM or VBQ statistical analysis. Whether and how MRI based brain morphometry and brain tissue property analysis will inform clinical decision making and clinical trial outcomes in spinal cord medicine remains to be determined.

## Introduction

Spinal cord injury (SCI) is a major cause for chronic disability that profoundly affects patients’ autonomy and quality of life. Despite the abundance of empirical evidence on the local effects of SCI along the spinal cord, our understanding of the concomitant changes in brain anatomy and function is still limited. Animal models of SCI showed controversial results ranging from extensive neuronal cell death in cortical areas (Hains et al., 2003) and the rubrospinal tract (Viscomi and Molinari, 2014) to lack of upper motoneuron degeneration or cell death of corticospinal neurons (Nielson *et al.*, 2010; Nielson *et al.*, 2011). The lack of in-depth knowledge about the impact of SCI on brain anatomy in humans highlights the need to provide *in vivo* analytic proof of concomitant structural changes that could inform clinical decision making in respect to treatment and prognosis.

Computational anatomy methods using magnetic resonance imaging (MRI) and mathematical algorithms to extract relevant brain features allow for statistical analysis of volume, shape and surface in three dimensional brain space (Ashburner *et al.*, 2003). One of the well-established methods – voxel-based morphometry (VBM), was used to monitor local grey matter volume changes following SCI to deliver conflicting results ranging from lack of SCI related brain anatomy changes (Crawley *et al.*, 2004) to evidence about profound sensorimotor cortex reorganization (Jurkiewicz *et al.*, 2006; Wrigley *et al.*, 2009a; Henderson *et al.*, 2011; Freund *et al.*, 2013). More recent reports demonstrate grey matter loss in non-motor areas including anterior cingulate gyrus, insula, orbitofrontal gyrus, prefrontal cortex and thalamus (Wrigley *et al.*, 2009b; Grabher *et al.*, 2015; Chen *et al.*, 2017). One of the potential reasons for the reported controversial findings is the fact that these studies pooled together patients with incomplete and complete SCI (Crawley *et al.*, 2004; Jurkiewicz *et al.*, 2006; Chen *et al.*, 2017) not taking into account potential impact of differences in the time span since injury (Grabher *et al.*, 2015). The non-quantitative character of the used T1-weighted MRI protocols represents another source for differences between studies. The computer-based estimation of regional volumes and cortical thickness from T1-weighted data is heavily dependent on the MR contrast, which is influenced by local histological tissue properties that give potentially rise to spurious morphological changes (Lorio *et al.*, 2016b).

Recent advances in quantitative MRI (qMRI) circumvent these limitations to provide quantitative maps indicative for myelin, iron and tissue free water content (Helms *et al.*, 2008; Draganski *et al.*, 2011; Lutti *et al.*, 2014). Investigations applying qMRI restricted to a set of regions-of-interest reported progressive volume loss in the internal capsule of SCI patients paralleled by myelin reduction at 12 months post-injury compared to baseline (Freund *et al.*, 2013). Using the same technique in the very same cohort, the authors observed also myelin reduction in thalamus, cerebellum and brainstem in the same period of time (Freund *et al.*, 2013; Grabher *et al.*, 2015). These combined morphometry and tissue property findings in the early injury phase contrast with the absence of volume differences when comparing sub-acute (duration <1 year) and chronic (duration >1 year) patients with complete motor SCI (Chen *et al.*, 2017).

Here we sought to address previous limitations in the field and investigate the sensitivity of quantitative brain tissue property MRI mapping to detect brain anatomy changes in a homogenous cohort of chronic complete SCI subjects We used the established voxel-based quantification (VBQ) and voxel-based morphometry to study potential differences in myelin, iron and free tissue water content between SCI subjects and healthy controls. We hypothesized that chronic complete SCI is associated with brain atrophy in the sensorimotor and non-sensorimotor system paralleled by specific alterations in myelin, iron and water content.

## Materials and Methods

### Study participants

All study related procedures were performed after obtaining informed consent according to protocols approved by the independent local ethics committee. We screened all patients admitted to the Spinal Cord Injury Center at Heidelberg University Hospital, Germany, for eligibility participating in the study. The main inclusion criterion was a spinal cord injury grade (American Spinal Injury Association Scale (AIS) grade A) that dated back at least 3 months before study participation. The control group was chosen with the intention to minimize age and gender differences between groups. Clinical scoring and grading were done according to the International Standards for Neurological Classification of Spinal Cord Injury (ISNCSCI) (Kirshblum *et al.*, 2011).

### MRI acquisition

All MRI data were acquired on a 3 Tesla scanner (Siemens Verio, Siemens Healthineers, Germany). The imaging protocol consisted of three whole-brain multi-echo 3D fast low angle shot (FLASH) acquisitions with predominantly magnetization transfer-weighted (MTw: TR/*α* = 24.5ms/6°), proton density-weighted (PDw: TR/*α* = 24.5ms/6°) and T1-weighted (T1w: 24.5ms/21°) contrast (Helms et al., 2009, 2008; Weiskopf et al., 2013). For each contrast we acquired multiple gradient echoes with minimum at 2.46ms and equidistant 2.46ms echo spacing. Per echo 176 sagittal partitions with 1mm isotropic voxel size (field of view and matrix size 256×240) and alternating readout polarity were acquired. The number of echoes was 7/8/8 for the MTw/PDw/T1w acquisitions to keep the TR value identical for all contrasts. We used parallel imaging along the phase-encoding direction (acceleration factor 2 with GRAPPA reconstruction) (Griswold et al., 2002) and partial Fourier (factor 6/8) along the partition direction.

### Map calculation

The R2*, MT, PD* and R1 quantitative maps were calculated as previously described (Draganski *et al.*, 2011). For map calculation we used in-house software running under SPM12 (Wellcome Trust Centre for Neuroimaging, London, UK; www.fil.ion.ucl.ac.uk/spm) and Matlab 7.11 (Mathworks, Sherborn, MA, USA). R2* maps were estimated from the regression of the log-signal of the eight PD-weighted echoes. MT and R1 maps were created using the MTw, PDw and T1w data averaged across all echoes (Helms *et al.*, 2008). All maps were corrected for local RF transmit field inhomogeneities using the inhomogeneity correction UNICORT algorithm in the framework of SPM (Weiskopf *et al.*, 2011).

### Voxel-based morphometry (VBM) and voxel-based quantification (VBQ)

For automated tissue classification in grey matter (GM), white matter (WM) and cerebrospinal fluid (CSF) we used the MT maps within SPMs “unified segmentation” approach (Ashburner and Friston, 2005) with default settings and enhanced tissue probability maps (Lorio *et al.*, 2016a) that provide optimal delineation of subcortical structures. Aiming at optimal anatomical precision, we estimated subject specific spatial registration parameters using the diffeomorphic algorithm based on exponentiated lie algebra – DARTEL (Ashburner, 2007). For VBM analysis, we scaled the probability maps with the corresponding Jacobian determinants to preserve the initial total amount of signal intensity. For VBQ analysis, we followed the same strategy by applying a weighted averaging procedure that ensures the preservation of the initial signal intensity of the MT, PD* and R2* parameter maps (Draganski *et al.*, 2011). The resulting maps were spatially smoothed using an isotropic Gaussian convolution kernel of 8 mm full-width-at-half-maximum.

### Spinal cord cross-sectional area (CSA) assessment

For CSA assessment we used the calculated T1-weighted images after contrast adjustment within snap-ITK (Yushkevich *et al.*, 2006). Delineation of spinal cord was performed by outlining the spinal cord circumference manually (AH) at the C2/3 level slice by slice in the axial plane yielding a total of 15 continuous slices. For an approximation of the mean cross-sectional area of the upper spinal cord the average of these 15 slices was used.

### Statistical analysis

We used parametric and nonparametric statistics from the software package JMP® – v12, for descriptive analysis of clinical data where deemed appropriate.

We created voxel-based parametric regression models with the group factor (SCI × healthy controls) as main predictor variable, and age, total intracranial volume (TIV – sum of GM, WM, and CSF tissue classes) and gender as additional variables (Barnes *et al.*, 2010) in an unpaired two-sample t-test design as implemented in SPM12. For the whole-brain analysis we reduced the search volume within brain’s GM or WM yielding 10 separate models (2 for VBM – GM-VBM and WM-VBM, 8 for VBQ – GM-PD*, GM-MT, GM-R1, GM-R2*, WM-PD*, WM-MT, WM-R1, WM-R2*). Parameter estimates and beta weights were estimated by appropriate one-sided t-contrast statements with corresponding statistical parametric T-maps. To control for multiple comparisons in this voxel-based analysis, family-wise error (FWE) correction methods using Random Field Theory were applied. The peak level height threshold for statistical significance was set at FWE p<0.05 with no cluster extent threshold.

## Results

### Population characteristics

MRI scans were obtained during a sampling period of 3.5 years. The main clinical characteristics of the patient population are summarized in table 1. We recruited 14 patients and 14 control subjects. The mean age in the control and patient group were 46±16 and 55±13 years (p=0.1147), respectively. The female male ratio was 3:11 in each group (p=1.0). The median time since SCI was 144 (14-568) months. Lesion severity in all patients was sensorimotor complete (AIS grade A). The TIV in the healthy control cohort was 1571±171ml (mean ± SD) and 1479±210ml (mean ± SD) – in SCI subjects (p=0.2169).

**Table 1:** Characteristics of SCI subjects. AIS (American Spinal Injury Association Impairment Scale), NLI (Neurological Level of Injury)

| ID | AIS | NLI | Age<br>(years) | Duration<br>(months) |
| --- | --- | --- | --- | --- |
| P1 | A | C7 | 66 | 531 |
| P2 | A | T7 | 62 | 519 |
| P3 | A | T10 | 32 | 206 |
| P4 | A | T5 | 59 | 122 |
| P5 | A | T3 | 55 | 114 |
| P17 | A | C5 | 21 | 7 |
| P19 | A | T5 | 59 | 4 |
| P20 | A | T8 | 65 | 4 |
| P21 | A | T6 | 61 | 66 |
| P22 | A | C6 | 53 | 16 |
| P25 | A | C3 | 52 | 385 |
| P30 | A | C6 | 65 | 568 |
| P32 | A | T9 | 62 | 567 |
| P34 | A | T5 | 58 | 165 |

### VBM and VBQ analysis

In the whole-brain VBM analysis we observed WM volume decreases in the dorsal part of the rostral cervical spinal cord in SCI subjects compared to healthy controls (mean difference 10±2 μl). There were no other significant grey or white matter brain volume differences (table 2; fig. 1).

**Table 2:** Voxel based morphometry significant findings (p<0.05 after FWE); WM = white matter; KE = cluster extent; T = t statistic; Z = z score statistic; coordinates = coordinates in Montreal Neurological Institute (MNI) space.

| tissue<br>map | contrast | regions | p FWE-corrected | K<sub>E</sub> | T | Z | coordinates |  |  |
| --- | --- | --- | --- | --- | --- | --- | --- | --- | --- |
|  |  |  |  |  |  |  | x | y | z |
| WM | SCI<Control | Medulla oblongata<br>(dorsal column) | 0.021 | 41 | 5.82 | 4.52 | -2 | -48 | -68 |

**Figure 1:**
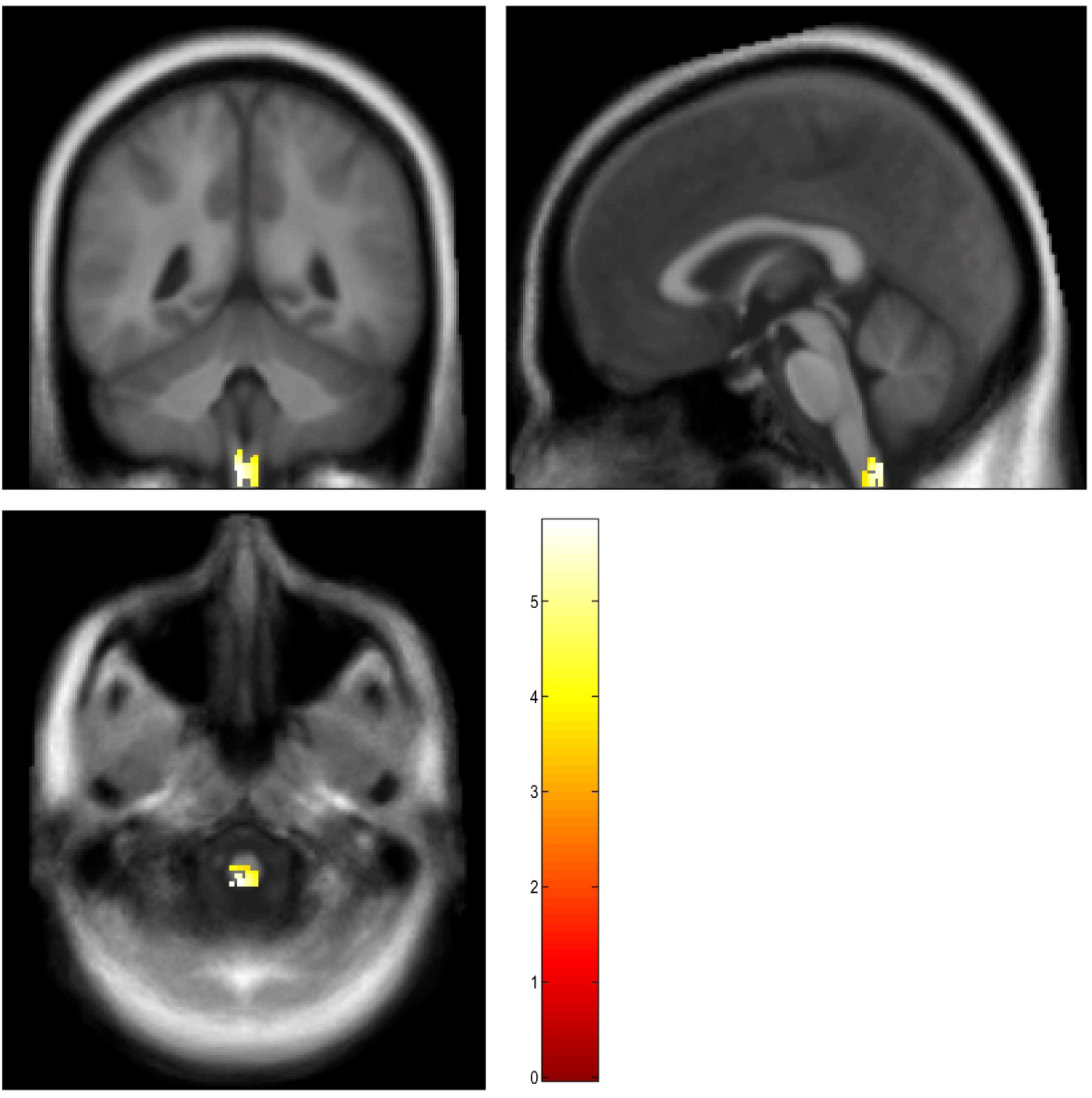
For illustration purposes representative section planes with significant voxels in VBM analysis at the alpha<0.001 uncorrected statistical threshold level are depicted. Color-coded voxels depict reduced WM volume in SCI subjects compared to healthy control individuals. The color scale represents T-values (height threshold: T=5.21, p=0.05 FWE)

The cross-sectional area (CSA) analysis at the cervical level revealed smaller CSA in SCI subjects compared to controls (60.2±8.1 mm^2^ versus 74.5±10.4, p<0.001). We found a positive correlation between the loss of cervical level CSA and WM volume at MNI -2 -48 -68 (r=0.662; p<0.001). CSA was not associated with any other WM and GM brain volume changes.

The VBQ analysis did not reveal any significant between-group differences.

## Discussion

In this study, we identified WM volume loss (approximately 10 μl) in the most rostral and dorsal part of the spinal cord in chronic sensorimotor complete SCI subjects. This volume loss correlated positively with spinal cord atrophy at the cervical level. In contrast to published results despite a reasonable sample size and a highly homogenous population from clinical and pathophysiological point of view, we were not able to find any morphometry or brain tissue property differences in SCI patients that reached the accepted levels of statistical significance.

The reduced volume in the dorsal region of the rostral spinal cord in SCI subjects most likely reflects Wallerian degeneration of large proprioceptive sensory axons, which has been confirmed histologically (Becerra *et al.*, 1995; Weber *et al.*, 2006). It is unlikely that the volume change at the rostral spinal cord was generated by a software algorithm induced imaging artifact due to false classification/registration (Bookstein, 2001). Quality inspection of the normalized images after application of the DARTEL algorithm did not indicate false classification or registration. The close positive correlation between spinal cord cross sectional area at the C2/3 level and the WM volume at the uppermost part of the spinal cord also supports the hypothesis of a true biological effect. Cord atrophy at the cervical and medulla oblongata level is a consistent finding after SCI, which is associated with more severe disability (Freund *et al.*, 2010; Freund *et al.*, 2011; Freund *et al.*, 2013). In other neurological diseases such as Parkinson’s disease volume changes were confirmed in similar regions (Jubault *et al.*, 2009).

WM volume reductions in the brain stem topographically related to the corticospinal tract and in the left cerebellar peduncle have been previously described in a more heterogeneous (more incomplete SCI subjects) and less chronic cohort of SCI subjects (Freund *et al.*, 2011; Freund *et al.*, 2013). However, volume changes in respective neuroanatomical regions were not reproducible in our more homogeneous chronic SCI cohort. Volume reductions in our cohort were located more caudal in the most rostrocaudal region of the cervical spinal cord consistent with the localization of the dorsal funiculus. Whether degenerative changes such as retrograde axon dieback or neuronal atrophy/cell death occur in corticospinal projections as suggested by VBM studies (Freund *et al.*, 2011; Freund *et al.*, 2013) is still debated. However, most recent preclinical studies indicate that respective alterations in the corticospinal tract – at least in rodents – cannot be observed (Nielson *et al.*, 2010; Nielson *et al.*, 2011). Remote changes in the brain related to neurogenesis or gliogenesis, which also could have produced volumetric changes in VBM studies (Killgore *et al.*, 2013), were not observed in animal models of SCI (Franz *et al.*, 2014). Post-mortem histological data from human SCI subjects are not available to either confirm or reject such changes.

Previous studies reported inconsistencies in respect to MRI based volumetric changes in the brain and affected regions in the brain (Crawley *et al.*, 2004; Jurkiewicz *et al.*, 2006; Wrigley *et al.*, 2009a; Freund *et al.*, 2011; Freund *et al.*, 2013; Hou *et al.*, 2014). In the current study, we were not able to attribute any VBM or VBQ brain or brainstem differences unambiguously to chronic sensorimotor complete SCI. The median time since injury was around 12 years in the analyzed cohort. Only one other study (Wrigley *et al.*, 2009a) analyzed a comparable SCI group in respect to injury severity and time since injury. However, our data do not support their finding of extensive reduced GM volumes in motor and non-motor regions of the brain related to SCI. We identified a correlation of brain volume reductions in respective regions only related to age (statistical significant covariate associated with smaller GM volumes; used as nuisance variable in our study), which has been consistently shown in previous studies not related to SCI (Raz *et al.*, 2007; Thompson and Apostolova, 2007; Draganski *et al.*, 2011). Lack of statistical power is also unlikely to explain the deviating findings. 14 SCI subjects were investigated in the present study, whereas 13 (Freund *et al.*, 2013), 10 (Freund *et al.*, 2011), 15 (Wrigley *et al.*, 2009a), and 17 (Jurkiewicz *et al.*, 2006) SCI subjects were enrolled in previous MRI studies.

In summary, our VBM and VBQ analyses in a highly homogenous group of chronic SCI subjects failed to detect main effects of chronic complete SCI on brain’s anatomy. These findings corroborate the absence of strong preclinical evidence of secondary neurodegenerative remote changes in the brain, yet confirm histological evidence in respect to remote changes in the spinal cord. At least with the methodology employed in the current study the application of MRI technology is not able to detect suitable and clinically meaningful markers in the brain, which help facilitate clinical decision making or enrich innovative clinical trial designs.

## Acknowledgements

This study was supported by grants from the Deutsche Forschungsgemeinschaft (SFB1158) to Norbert Weidner and rom the Medical Faculty Heidelberg to Andreas Hug. We thank Bogdan Draganski from the Department of Clinical Neurosciences Lausanne University Hospital, University of Lausanne, Switzerland, for his expert advice.

## References

Ashburner J. A fast diffeomorphic image registration algorithm. Neuroimage 2007; 38(1): 95–113.

Ashburner J, Csernansky JG, Davatzikos C, Fox NC, Frisoni GB, Thompson PM. Computer-assisted imaging to assess brain structure in healthy and diseased brains. Lancet Neurol 2003; 2(2): 79–88.

Ashburner J, Friston KJ. Unified segmentation. Neuroimage 2005; 26(3): 839–51.

Barnes J, Ridgway GR, Bartlett J, Henley SM, Lehmann M, Hobbs N, et al. Head size, age and gender adjustment in MRI studies: a necessary nuisance? Neuroimage 2010; 53(4): 1244–55.

Becerra JL, Puckett WR, Hiester ED, Quencer RM, Marcillo AE, Post MJ, et al. MR-pathologic comparisons of wallerian degeneration in spinal cord injury. AJNR Am J Neuroradiol 1995; 16(1): 125–33.

Bookstein FL. “Voxel-based morphometry” should not be used with imperfectly registered images. Neuroimage 2001; 14(6): 1454–62.

Chen Q, Zheng W, Chen X, Wan L, Qin W, Qi Z, et al. Brain Gray Matter Atrophy after Spinal Cord Injury: A Voxel-Based Morphometry Study. Front Hum Neurosci 2017; 11: 211.

Crawley AP, Jurkiewicz MT, Yim A, Heyn S, Verrier MC, Fehlings MG, et al. Absence of localized grey matter volume changes in the motor cortex following spinal cord injury. Brain Res 2004; 1028(1): 19–25.

Draganski B, Ashburner J, Hutton C, Kherif F, Frackowiak RS, Helms G, et al. Regional specificity of MRI contrast parameter changes in normal ageing revealed by voxel-based quantification (VBQ). Neuroimage 2011; 55(4): 1423–34.

Franz S, Ciatipis M, Pfeifer K, Kierdorf B, Sandner B, Bogdahn U, et al. Thoracic rat spinal cord contusion injury induces remote spinal gliogenesis but not neurogenesis or gliogenesis in the brain. PLoS One 2014; 9(7): e102896.

Freund P, Weiskopf N, Ashburner J, Wolf K, Sutter R, Altmann DR, et al. MRI investigation of the sensorimotor cortex and the corticospinal tract after acute spinal cord injury: a prospective longitudinal study. Lancet Neurol 2013; 12(9): 873–81.

Freund P, Weiskopf N, Ward NS, Hutton C, Gall A, Ciccarelli O, et al. Disability, atrophy and cortical reorganization following spinal cord injury. Brain 2011; 134(Pt 6): 1610–22.

Freund PA, Dalton C, Wheeler-Kingshott CA, Glensman J, Bradbury D, Thompson AJ, et al. Method for simultaneous voxel-based morphometry of the brain and cervical spinal cord area measurements using 3D-MDEFT. J Magn Reson Imaging 2010; 32(5): 1242–7.

Grabher P, Callaghan MF, Ashburner J, Weiskopf N, Thompson AJ, Curt A, et al. Tracking sensory system atrophy and outcome prediction in spinal cord injury. Ann Neurol 2015; 78(5): 751–61.

Hains BC, Black JA, Waxman SG. Primary cortical motor neurons undergo apoptosis after axotomizing spinal cord injury. J Comp Neurol 2003; 462(3): 328–41.

Helms G, Dathe H, Kallenberg K, Dechent P. High-resolution maps of magnetization transfer with inherent correction for RF inhomogeneity and T1 relaxation obtained from 3D FLASH MRI. Magn Reson Med 2008; 60(6): 1396–407.

Henderson LA, Gustin SM, Macey PM, Wrigley PJ, Siddall PJ. Functional reorganization of the brain in humans following spinal cord injury: evidence for underlying changes in cortical anatomy. The Journal of neuroscience : the official journal of the Society for Neuroscience 2011; 31(7): 2630–7.

Hou JM, Yan RB, Xiang ZM, Zhang H, Liu J, Wu YT, et al. Brain sensorimotor system atrophy during the early stage of spinal cord injury in humans. Neuroscience 2014; 266: 208–15.

Jubault T, Brambati SM, Degroot C, Kullmann B, Strafella AP, Lafontaine AL, et al. Regional brain stem atrophy in idiopathic Parkinson’s disease detected by anatomical MRI. PLoS One 2009; 4(12): e8247.

Jurkiewicz MT, Crawley AP, Verrier MC, Fehlings MG, Mikulis DJ. Somatosensory cortical atrophy after spinal cord injury: a voxel-based morphometry study. Neurology 2006; 66(5): 762–4.

Killgore WD, Olson EA, Weber M. Physical exercise habits correlate with gray matter volume of the hippocampus in healthy adult humans. Sci Rep 2013; 3: 3457.

Kirshblum SC, Waring W, Biering-Sorensen F, Burns SP, Johansen M, Schmidt-Read M, et al. Reference for the 2011 revision of the International Standards for Neurological Classification of Spinal Cord Injury. The journal of spinal cord medicine 2011; 34(6): 547–54.

Lorio S, Fresard S, Adaszewski S, Kherif F, Chowdhury R, Frackowiak RS, et al. New tissue priors for improved automated classification of subcortical brain structures on MRI. Neuroimage 2016a; 130: 157–66.

Lorio S, Kherif F, Ruef A, Melie-Garcia L, Frackowiak R, Ashburner J, et al. Neurobiological origin of spurious brain morphological changes: A quantitative MRI study. Hum Brain Mapp 2016b; 37(5): 1801–15.

Lutti A, Dick F, Sereno MI, Weiskopf N. Using high-resolution quantitative mapping of R1 as an index of cortical myelination. Neuroimage 2014; 93 Pt 2: 176–88.

Nielson JL, Sears-Kraxberger I, Strong MK, Wong JK, Willenberg R, Steward O. Unexpected survival of neurons of origin of the pyramidal tract after spinal cord injury. The Journal of neuroscience : the official journal of the Society for Neuroscience 2010; 30(34): 11516–28.

Nielson JL, Strong MK, Steward O. A reassessment of whether cortical motor neurons die following spinal cord injury. J Comp Neurol 2011; 519(14): 2852–69.

Raz N, Rodrigue KM, Haacke EM. Brain aging and its modifiers: insights from in vivo neuromorphometry and susceptibility weighted imaging. Ann N Y Acad Sci 2007; 1097: 84–93.

Thompson PM, Apostolova LG. Computational anatomical methods as applied to ageing and dementia. Br J Radiol 2007; 80 Spec No 2: S78–91.

Viscomi MT, Molinari M. Remote neurodegeneration: multiple actors for one play. Mol Neurobiol 2014; 50(2): 368–89.

Weber T, Vroemen M, Behr V, Neuberger T, Jakob P, Haase A, et al. In Vivo High-Resolution MR Imaging of Neuropathologic Changes in the Injured Rat Spinal Cord. 2006; 27(3): 598–604.

Weiskopf N, Lutti A, Helms G, Novak M, Ashburner J, Hutton C. Unified segmentation based correction of R1 brain maps for RF transmit field inhomogeneities (UNICORT). Neuroimage 2011; 54(3): 2116–24.

Wrigley PJ, Gustin SM, Macey PM, Nash PG, Gandevia SC, Macefield VG, et al. Anatomical changes in human motor cortex and motor pathways following complete thoracic spinal cord injury. Cereb Cortex 2009a; 19(1): 224–32.

Wrigley PJ, Press SR, Gustin SM, Macefield VG, Gandevia SC, Cousins MJ, et al. Neuropathic pain and primary somatosensory cortex reorganization following spinal cord injury. Pain 2009b; 141(1-2): 52–9.

Yushkevich PA, Piven J, Hazlett HC, Smith RG, Ho S, Gee JC, et al. User-guided 3D active contour segmentation of anatomical structures: significantly improved efficiency and reliability. Neuroimage 2006; 31(3): 1116–28.

